# Not dasycladalean alga, but an Odyssey of the earliest Phanerozoic animal reef-builders

**DOI:** 10.1101/2023.12.07.570709

**Authors:** Aihua Yang, Cui Luo, Jian Han, Andrey Yu. Zhuravlev, Joachim Reitner, Haijing Sun, Han Zeng, Fangchen Zhao, Shixue Hu

## Abstract

The compacted macrofossil *Protomelission*? sp. from the early Cambrian Xiaoshiba Lagerstätte was recently ascribed to early dasycladalean green algae and used to disprove the bryozoan affinity of coeval phosphatized microfossils, which made the puzzling question whether the bryozoans originated in early Cambrian pending again. Our new analyses of multiple specimens which are conspecific with *Protomelission*? from the Chengjiang Lagerstätte indicate that they are not dasycladaleans but one of the three groups of archaeocyath-like sponges that atypically inhabited siliciclastic substrates. All the archaeocyath-like fossils share the same preservation mode and exhibit archaeocyath-type external skeletal features. Particularly, the *Protomellision*?-like fossils preserve structures indicative of archaeocyath aquiferous system and ontogeny. They represent the first recognized one-walled archaeocyath sponges in South China and evidence the niche expansion of archaeocyaths on their way of global radiation from Siberia, 518 million years ago. The origin of the bryozoans remains a mystery.

---

The Cambrian explosion is the most significant evolutionary event in Earth history in terms of setting up stem- and even crown-groups of almost all major metazoan phyla^1^ and the Phanerozoic-type, animal-dominated ecosystem^2, 3^. A current debate is about possible Cambrian origin of the phylum Bryozoan, whose distinct oldest fossil record was traced back to the early Ordovician only^4^. Recently, the phosphatized microfossils *Protomelission gatehousei* from the Wirrealpa Limestone, South Australia, and the Xihaoping Member, South China were interpreted as early Cambrian bryozoans^5^. On the contrary, a compacted macrofossil *Protomelission*? sp. from the Xiaoshiba Lagerstätte mudstone was ascribed to dasycladalean green algae, allowing the authors to suggest algal affinities of *P. gatehousei*^6^.

The identification of *Protomelission*? sp. and *P. gatehousei* as dasycladalean algae suffers from several inconsistencies. Firstly, the Xiaoshiba fossil differs drastically from *P. gatehousei* both in size and construction. *P. gatehousei* is a microscopic (2.2 mm in height or less) three-dimensional, phosphatized bifoliate lamellar plate with an elliptical holdfast^5^. Chambers with large openings are arranged back-to-back on the two sides of this plate (Figs. 2 and 3 in Zhang *et al*.)^5^. By contrast, the highly compressed *Protomelission*? sp. is a two-order larger (108–160 mm in height) subcylindrical fossil with a single-layered wall^6^. A preservational selectivity in favor of smaller phosphatized fragments in the Wirrealpa Limestone is irrelevant to these discrepancies because numbers of sizable and fragile phosphatic fossils have been described from this unit^7^. Secondly, *Protomelission*? sp. itself is also not necessarily a dasycladalean alga. Of about 180 mostly extinct genera of dasycladalean algae, none has a morphology resembling *Protomelission*? sp.^8^. The principal features of these algae are rooted in their siphonous organization, which is expressed in a monopodial thallus with laterals of several orders^8,9^.

Here, based on well-preserved fossils from the Chengjiang Lagerstätte (Cambrian Stage 3), we argue that *Protomelission*? sp. from the Xiaoshiba Lagerstätte belongs to one of the archaeocyath taxa that are preserved in early Cambrian fine siliciclastic taphonomic windows. Archaeocyaths were calcareous cup-like aspiculate sponges, which built the earliest Cambrian reefs with some other organisms and strictly occupied carbonate substrates^10^. This is the first discovery of their niche expansion to the siliciclastic settings.

## Results

### Morphology and preservation of the archaeocyath-like fossils

Three sets of specimens were recognized as potential archaeocyaths from the Chengjiang biota (see Methods), here named archaeocyath-like species 1, 2, and 3 (ASP1, 2, and 3). They are preserved on the bedding plane as narrow conical to subcylindrical flattened bodies up to more than 80 mm high and 10 mm wide at the top (Supplementary Table 1). These fossils, probably, are skeletons with somewhat stiffness because they do not display twisted or folded preservation typical of fleshy algae.

Six ASP1 specimens are nearly indistinguishable from the reported *Protomelission*? sp. in appearance (Fig. 1a). These fossils are 4–9 mm wide and can be over 63 mm long (Supplementaty Table 1). In general, they are characterized by quincuncially or diagonally arranged bulges on the surface (Fig. 1a), comparable with those of *Protomelission*? sp. (Figs. 1–2 in ref.^6^). Upon a closer view, the bulges in the upper part of the fossil are tiny, downward-orienting tubular canals buried in the host mudstone (Fig. 1d). While toward the lower part of the organism, the canals shorten and grade into raised margins surrounding hexagonal to rounded depressions (Fig. 1f–i). In a specimen, which represents a topmost fragment of the organism, the canals are so well developed that they impart the fossil a pattern of elongated rhombic grids on the surface (Fig. 1j). The canals are non-or only weakly mineralized and show features of soft compaction (arrows in Fig. 1k). The axial rows of canals are most strongly flattened because they are underlain by the thickest part of the lenticular sedimentary infill in the inner cavity (Fig. 1j, k).

**Fig. 1.**
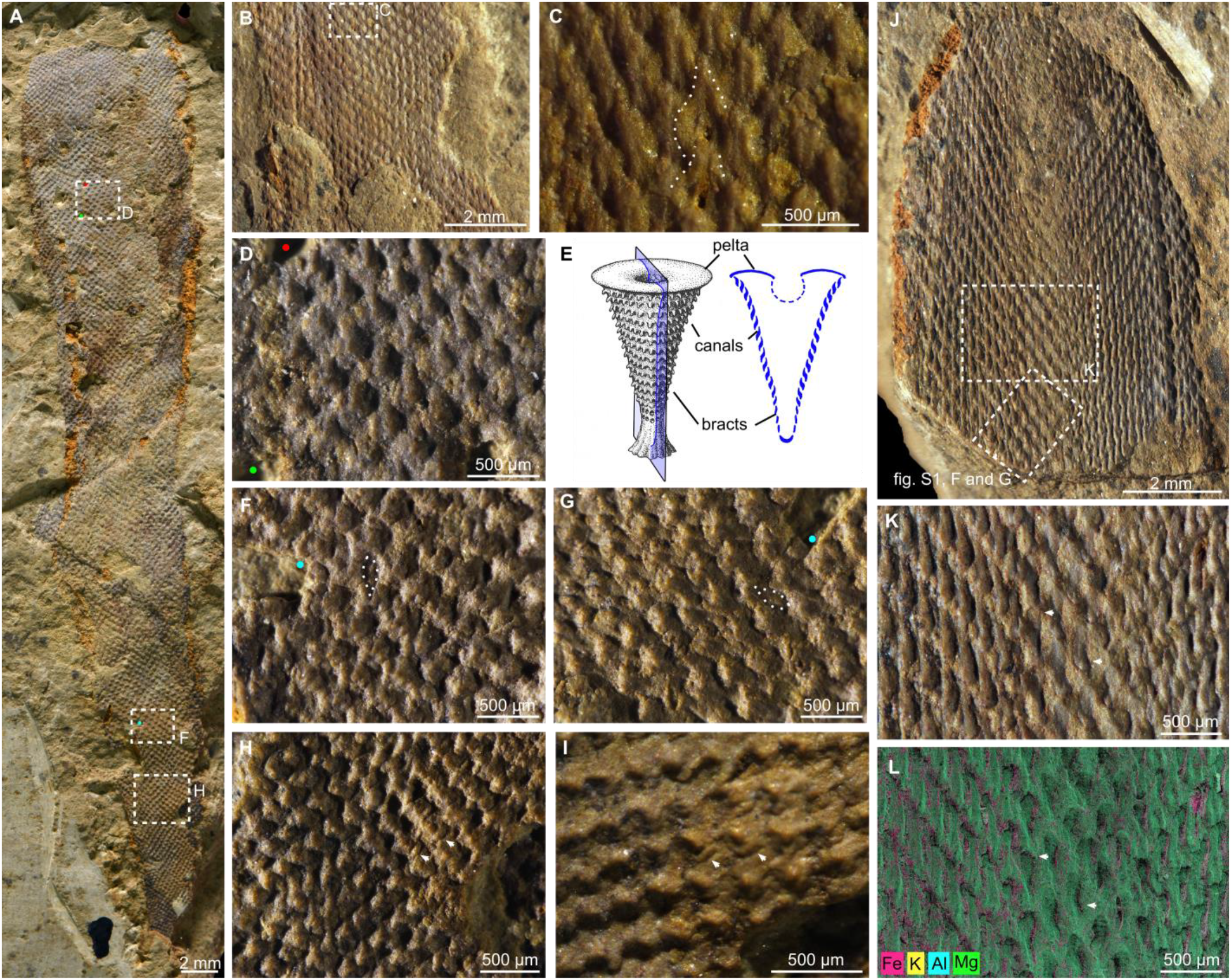
Morphology and preservation of possible archaeocyath ASP1 from the Chengjiang Lagerstätte, South China. **a** General view of AC-03-JS, areas in rectangles enlarged in **d, f, h**, respectively. **b** AC-02-JSa, showing the inner cavity within the single wall partly filled by sediments, rectangle enlarged in **c. c** Close view of the internal surface of the wall, showing tiny pores and straight furrows (emphasized with dash lines), which probably represent the proximal termination of the tubular canal. **d, f–i** Close view of different parts of the outer surface of AC-03-JS. **g** and **i** represent a rotated view of **f** and **h**, respectively; dots, dashed lines, and arrows indicate corresponding areas in paired images. **e** Reconstruction (left) of one-walled archaeocyath *Propriolynthus* and its axial longitudinal section (right) adapted after Debrenne et al.^11^. **j** AC-08-JSa showing the top of the specimen with the area in rectangle enlarged in **k. k** Flattened canals (arrows). **l** Elemental mapping of the area in **k**, arrows point to the same features as in **k**.

**Fig. 2.**
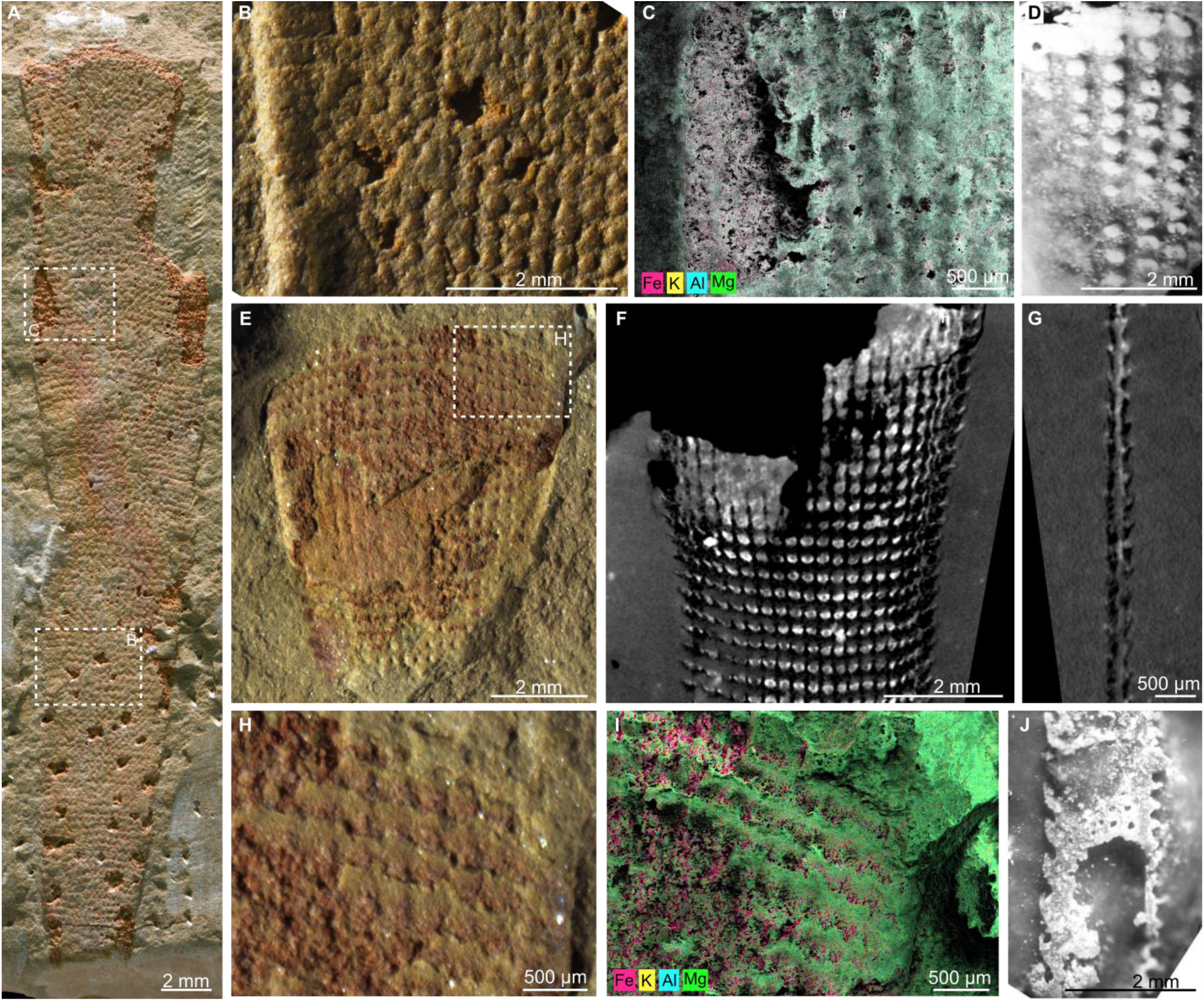
Morphology and preservation of possible archaeocyaths ASP2 and ASP3 from the Chengjiang Lagerstätte and similar features of archaeocyaths from the lower Cambrian of South Australia. **a** AC-10-SKa, a specimen of ASP2, with areas in rectangles enlarged in **b, c. b** Close view of a part of **a**, showing bulges on the outer surface. **c** Elemental mapping of a part of **a. d** Simple tumuli on the outer wall of *Ethmocoscinus papillipora*, PU 86679. **e** AC-07-SKa, a specimen of ASP3. **f, g** AC-07-SKb, the counterpart of **e**, in micro-CT, showing three-dimensionally preserved bracts. **h, i** A close view of ASP3 bracts (**h**) and its elemental mapping (**i**) showing preservation of biogenic structures in clay minerals. **j** Outer wall scales (right) of *Sigmocoscinus sigma*, USNM PU86686.

This compound wall is single-layered, with siliciclastic mud infilled its inner cavity (Figs. 1b and Supplementary Fig. 1a, b). The internal surface of the wall has a reverse relief compared with the outer one (Figs. 1b, c and Supplementary Fig. 1a–e, h, i). Tiny pores are visible on the internal surface, corresponding to the proximal and topmost terminations of the canals (Figs. 1c and Supplementary Fig. 1c–e). The hexagonal to rounded depressions on the outer surface are related either to the distal tips of these canals, which are protected by bracts (Fig. 1f–i), or are resulted from compression of these elongated canals (Figs. 1k and Supplementary Fig. 1f, g).

With all these characters, ASP1 can be assigned to one-walled archaeocyaths, represented by the order Monocyathida^12^. The upward transition from bracts/scales to canals on the wall is comparable with the ontogenetic model of *Propriolynthus*^11^ (Fig. 1e). Numbers of monocyathides possess peltae, which are slightly convex plates developed as a continuation of the wall at the top of the cup and covering the inner cavity to restrict its opening^11,13^ (Fig. 1e). Accordingly, no large opening of the inner cavity is detected in ASP1.

ASP2 and ASP3 are both characterized by an orthogonal arrangement of structural elements but are distinguished from each other by the wall features. The former exhibits small hollow domes organized in regular longitudinal and transverse rows (Fig. 2a–c). The arrangement of these domes corresponds to the orthogonal grids in the underlying skeleton (Fig. 2a, b). This outer wall structure is comparable with tumuli in archaeocyaths like *Tumuliolynthus* and *Ethmocoscinus*^11,13^ (Fig. 2d). ASP3 bears transverse external narrow plate-like structures on the porous wall (Fig. 2e–i), which can be described as upwardly projecting planar annuli or scales in terms applicable to archaeocyathan morphology^11,13^ (Fig. 2j). Alike outer wall structure is known from multiple archaeocyath taxa, including the one-walled archaeocyath *Robertiolynthus*^12^. The internal structures in ASP2 and ASP3 are not finely preserved as those in ASP1 due to compaction and replacement by iron oxides.

The authentic structures on the outer surface of all the ASP fossils are delicately preserved in clay minerals, while iron oxides are concentrated around the veneer of clay minerals (Figs. 1k, l, 2c, i, and Supplementary Fig. 1c–i). This is also evident in micro-CT images, where dark bracts are surrounded by a brighter host matrix. The latter has a higher density due to elevated metal content (Fig. 2f, g).

## Discussion

### Comparisons among new fossils, *Protomelission*? sp., archaeocyaths, and dasycladaleans

As emphasized above, ASP1 is almost identical to *Protomelission*? sp. in morphology except for the absence of flanges and budding (cf. Fig. 2f, g in ref. ^6^ and ASP1 described here). Their preservation is likewise comparable. Yang *et al*.^6^ stated that “each module is enclosed by a thin but robust layer associated with elevated concentrations of iron and phosphorous”. This is in accordance with our observation that the spaces between canals are filled by iron-rich deposites (Fig. 1k, l and Supplementary Fig. 1d–g). Similarly, the elemental maps in the Extended Data Fig. 1 of Yang *et al*.^6^ show that elevated concentrations of iron correspond to visible black metal dyeing and to the red iron oxide crust. The biogenic structures, such as the flanges, are preserved with clay minerals. The elevation of phosphorus in *Protomelission*? sp. is doubtful. In Extended Data Fig. 1 of Yang et al.^6^, phosphorus is universal in the host matrix. Phosphorus is also present in all the Chengjiang fossils studied here but only with a very low concentration (0.2–0.3 wt%) (Supplementary Fig. 2h, p, x). Unfortunately, Yang *et al*.^6^ did not provide data on the relative concentrations of elements.

Irrespectively of their absence in our specimens, flanges are not a typical feature of dasycladaleans^8,9^ (Fig. 3b), except for the heavily calcified middle Cambrian problematic fossil *Amgaella* from the Siberian Platform^9^ (Fig. 3f, g). However, morphological interpretation of the middle-upper Cambrian family Seletonellaceae, which includes *Amgaella*, is based on highly imaginary reconstructions^14,15^. The dasycladalean or general algal affinity of this taxon is still pending^16^. The “hair” attached to the tip of laterals in some living dasycladaleans (Fig. 3c) is entirely different from the flanges described in Yang *et al*.^6^ since the former are extremely slender and soft^9^. Besides, the budding described in Yang *et al*.^6^ discords the dasycladalean interpretation but fits the array of archaeocyath developmental features^17^. The latter includes external budding similar with that of *Protomelission*? sp. Dasycladalean green algae are unicellular organisms characterized by a monopodial thallus, which is differentiated into an upright cylindrical central axis and whorls of appendage-like laterals (Fig. 3b, c). The central axis is usually unbranched and rarely dichotomously branched^9^. Moreover, the tubular canals in ASP1 are downward pointing. This growth habit is unknown in any algal branches, but fits to a sponge model: the downward pointing inhalant pores are less likely clogged by detritus in siliciclastic environments^13^.

**Fig. 3.**
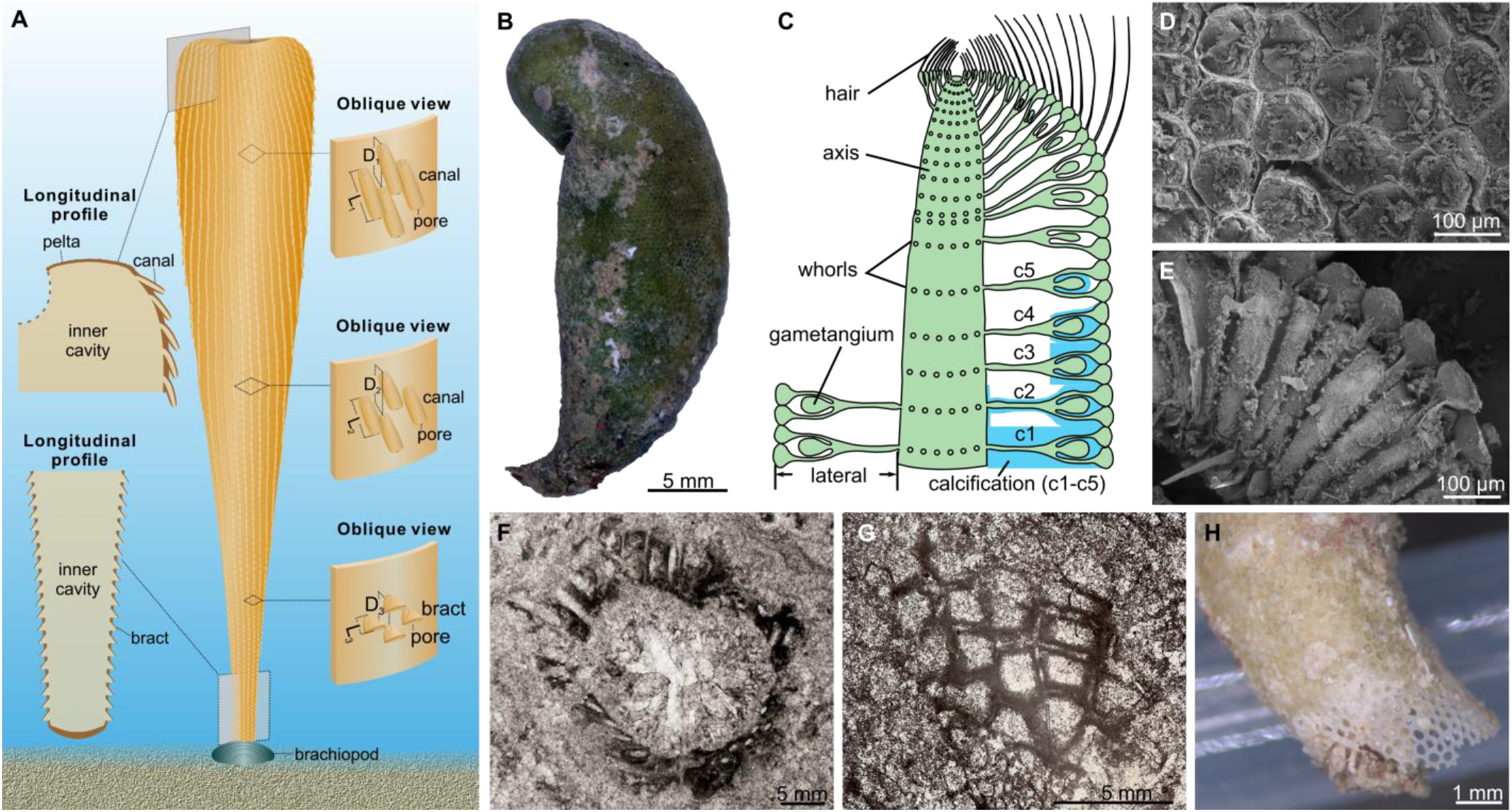
Comparisons among ASP1, dasycladalean algae, and *Amgaella*. **a** reconstruction of ASP1 based on this study. Annotations: D, distances between pores, note that D_1_ > D_2_ > D_3_; L, length of the canals and bracts, note that L_1_ > L_2_ > L_3_. **b** A complete thallus of the living dasycladalean alga *Neomeris annulate* from the intertidal zone of Bermuda. **c** Drawing of the longitudinal section of *Neomeris* modified after Berger and Kaever^9^. Color code: green, thallus; blue, possible positions and intensities of calcification (c1–c5). **d, e** SEM observation of *N. annulata*, showing its surface covered by the inflated tips of the laterals (**d**), and aragonite deposition in the interspace among laterals (**e**). **f, g** Transverse (**f**) and tangential (**g**) sections of the middle Cambrian problematic fossil *Amgaella*. (**h**) The aragonitic frame of *Neomeris* without organic tissue.

The preservation mode raises another question about the dasycladalean interpretation of *Protomelission*? sp. Dasycladaleans are preserved either as carbonaceous compressions or as calcified body fossils if, when alive, they have precipitated aragonite in the mucilage outside of the cell wall^9,18^. These mineral skeletons are essentially an external mould of the thalli (Fig. 3c, e, h). In non-calcified dasycladalean fossils, the thalli are often composed of bush-like, branching filaments and sometimes bear reproductive structures such as cysts and gametophores^19–25^. These compressions are enriched with carbon and sulfur but are depleted with iron and phosphorus^26,27^. Although some dasycladaleans form a so-called “pseudocortex” by the inflation of the distal tip of the laterals (Fig. 3c–e), the above-mentioned construction of the thallus determines that any expectable cup-like fossils of dasycladaleans are restricted to their mineralized ‘exoskeletons’.

So far, archaeocyath seems to be the best model for the affinity of ASP1 and *Protomelission*? sp. Apart from archaeocyaths, a few other Cambrian skeletal fossils display a similar conical shape with evenly distributed angular orifices on the external surface. These are *Tunkia* from southern Australia and *Lenaella* from the Siberian Platform; both possess an external ornament comprising two intersecting sets of diagonal ribs, together bounding rhomboid pits. Whereas their internal surface is smooth and aporous, and the aperture lacks a cover^28^. Furthermore, the gradual upward transition from bracts to canals in ASP1 fits the archaeocyathan skeletal ontogeny but not to features of either bryozoans or dasycladaleans, which do not display any ontogenetic changes in their skeletons.

### Niche expansion and paleogeographic migration of archaeocyaths

Archaeocyaths, now broadly accepted as an extinct class of the phylum Porifera^12,13^, were the earliest reef-building metazoans in the Phanerozoic^30^. Although they have reached a biodiversity of over 300 genera and ca. 1500 species before the end of the Cambrian Age 4, their habitats are generally limited to tropical normal-marine carbonate depositional environments. The most significant deviation from this habitat preference known before are archaeocyath-microbe carbonate bioherms developed on siliciclastic substrates^13,31,32^. Because of this narrow habitat preference, the global dispersal of archaeocyath taxa were believed to be controlled by the availability of adjacent suitable areas for archaeocyaths to live^13^. This model worked well in interpreting the provincialism of archaeocyath fauna around the world. The fossils described here are the first example of ahermatypic and non-calcified archaeocyaths occurring in siliciclastic environments. Notably, ASP1 and *Protomelission*? sp. belongs to the order Monocyathida, which has never been observed in South China^11,33^, although their occurrence in this terrane is expected based on published data of global distribution of these one-walled archaeocyaths over time (Fig. 4). This discovery of monocyathides in Chengjiang and Xiaoshiba Lagerstätten not only closes the missing link in the spreading route of monocyathides (Fig. 4), but also reveals a new means of their global migration. That is, spreading by adapting to siliciclastic depositional environments (Fig. 4). A few animal groups promoted niche expansion during their evolution, leading to phenotypic diversification (for instance, ref. ^34^). Similarly, archaeocyaths were able to expand their common niche to siliciclastic environments by releasing from their calcareous ‘casual clothes’, and this allowed them to evolve into more familiar aspiculate sponges^35^.

**Fig. 4.**
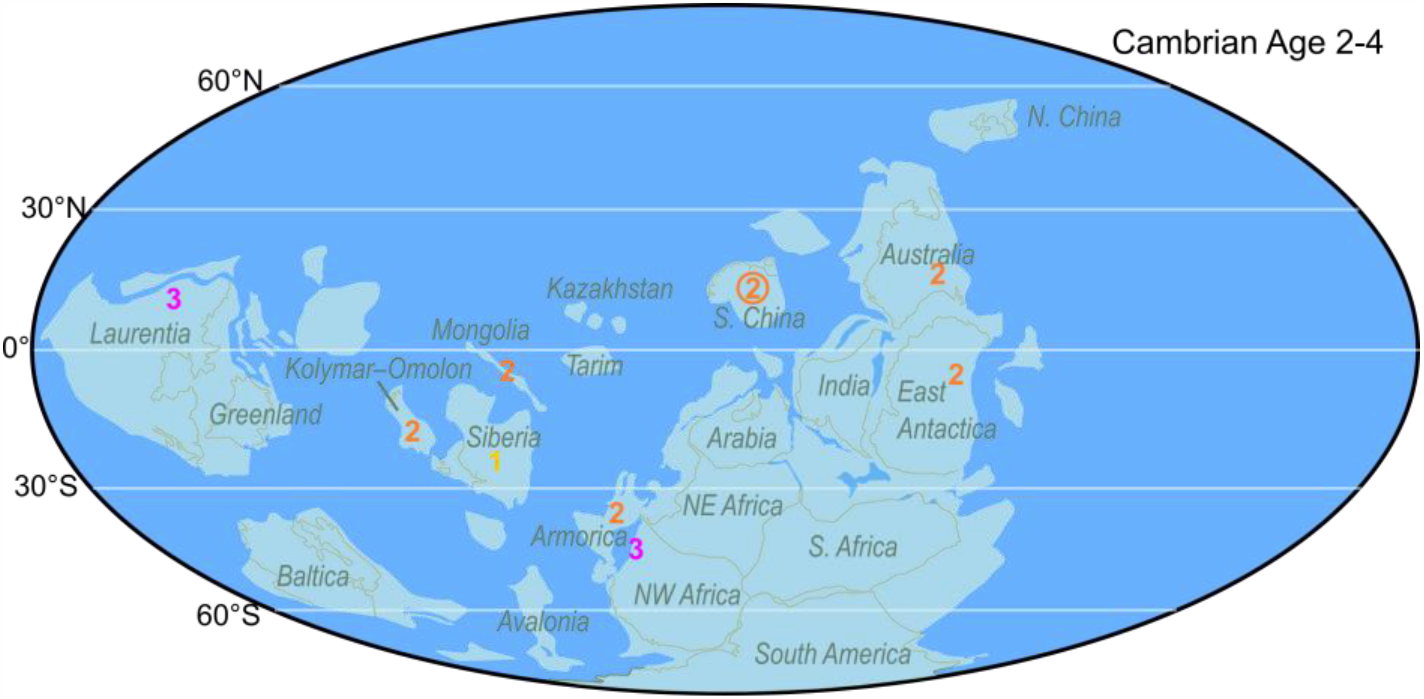
Radiation of monocyathide archaeocyaths during Cambrian Age 2–4. Paleogeographic map modified after ref. ^29^. Distribution of monocyathides visualized after ref. ^11^. Numbers indicate the age of the earliest appearance of monocyathides in the relevant localities: 1, Tommotian (late Age 2); 2, Atdabanian (early to middle Age 3); 3, Botoman (late Age 3 to early Age 4). The number with circle indicates the fossils described in this work.

## Methods

### Material

Specimens of ASP1–3 were discovered in the Lagerstätte-bearing Maotianshan Shale Member of the Yu’anshan Formation from the Shankoucun section in the Anning area and the Jianshan section in the Haikou area, Yunnan Province, South China. The Chengjiang Lagerstätte is restricted to the middle-upper *Eoredlichia-Wudingaspis* trilobite Zone of Cambrian Stage 3, Series 2^25^.

Every specimen is derived from event beds of the Maotianshan Shale Member and represents a toppled and compressed narrow conical skeleton with its longitudinal axis oriented parallel to bedding and highlighted by a rusty stain. Each specimen is easily split along its external surface into a part and two counterparts: one from each side of a flattened lenticular fossil (the part itself).

The illustrated fossil specimens from the Chengjiang Lagersätte are reposited in the Northwest University, Xi’an (NWU) and the Nanjing Institute of Geology and Palaeontology, Chinese Academy of Science (NIGPAS), China. Specimens of *Neomeris* are archived at the University of Göttingen. *Amgaella* is housed in the Siberian Scientific-Research Institute of Geology, Geophysics and Natural Resources, Novosibirsk. The two Australian archaeocyaths are archived at the National Museum of Natural History, Smithsonian Institution, Washington, D.C. (USNM) and the Princeton University, Princeton, New Jersey (PU).

### Stereomicroscopy

Microscopic study and photographing of the fossils were achieved using a Nikon SMZ18 stereomicroscope coupled with a Nikon DSRi2 camera at the Nanjing Institute of Geology and Palaeontology, Chinese Academy of Sciences (NIGPAS).

### Scanning electron microscopy (SEM) and energy-dispersive X-ray spectroscopy (EDS)

Elemental mapping of uncoated and unpolished samples including fossil and adjacent matrix was performed at NIGPAS, with Oxford Ultim Max energy-dispersive X-ray spectroscopy coupled to a TESCAN MAIA 3 GMU scanning electron microscope under an electron beam accelerating voltage of 15 kV.

### X-ray computed tomography

X-ray computed tomography (CT) was carried out at the Micro-CT lab (NIGPAS) using a Zeiss Xradia 520 Versa instrument with 50 kV operating voltage of the X-ray tube and with a thin filter (LE1) to avoid beam-hardening biases for a non-destructive 3D-reconstruction including an animation of a cup preserving the most features. For each scan, 3000 equiangular projections over 360ºwere obtained with an exposure time of 10 s per projection; the resulting volume data were processed using VG Studio Max 2.0 software.

## Data availability

Data supporting the findings of this study are available within the paper and its supplementary files. The source data and parameters of the CT scans can be accessed at https://… (original data will be uploaded upon the acceptance of the manuscript).

## Acknowledgements

We thank Y. Fang and S.-p. Wu for assistance SEM and EDS analyses and micro-CT data processing, respectively. We are also grateful to Prof. M. Zhu and Prof. Z. Yin for proofreading and suggestions on the text. This study was supported by the National Key Research and Development Program of China (2022YFF0800100) and National Natural Science Foundation of China (41872011, 41921002, 42072006, 41972016).

## Author contributions

A.-h.Y., A.Y.Z., J.H. and F.-c.Z. designed the project. The Chengjiang fossil materials were contributed by A.-h.Y., J.H., F.-c.Z., S.-x.H. and J.R.; *Neomeris* photos by J.R.; photos of *Amgaella* and Australian archaeocyaths by A.Y.Z. C.L. carried out the microscopy observation and photography, SEM and EDS analyses, and assembled all data and figure plates. A.-h.Y. organized the micro-CT scan. H.Z. made DSLR photography of all specimens. H.-j.S. and C.L. finished the reconstruction drawing of ASP1. With the input from A.-h.Y., C.L. and A.Y.Z. wrote the first version of the manuscript. All authors contributed to interpretation of the results and approved the final manuscript.

## Competing interests

The authors declare no competing interests.

## Additional information

**Supplementary information** The online version contains supplementary material available at https://.

## Notes

### Competing Interest Statement

The authors have declared no competing interest.

## References

1. Erwin, D. H., Laflamme, M. S., M. Tweedt, Sperling, E. A., Pisani, D., Peterson, K. J., The Cambrian Conundrum: Early divergence and later ecological success in the early history of animals. Science 334, 1091–1097 (2011).

2. Dunne, J. A., Williams, R. J., Martinez, N. D., Wood, R. A. Erwin, D. H., Compilation and network analyses of Cambrian food webs. PLOS Biology 6, e102 (2008).

3. Mángano, M. G., Buatois, L. A., The rise and early evolution of animals: where do we stand from a trace-fossil perspective? Interface Focus 10, 20190103 (2020).

4. Taylor, P. D., Bryozoan Paleobiology (Wiley Blackwell, London, UK, 2020).

5. Zhang, Z., Zhang, Z., Ma, J. Taylor, P. D., Strotz, L. C., Jacquet, S. M., Skovsted, C. B., Chen, F., Han Brock, J. G. A., Fossil evidence unveils an early Cambrian origin for Bryozoa. Nature 599, 251–255 (2021).

6. Yang, J., Lan, T., Zhang, X., Smith, M. R., Protomelission is an early dasyclad alga and not a Cambrian bryozoan. Nature 615, 468–471 (2023).

7. Skovsted, C. B., Brock, G. A., Topper, T. P., Paterson, J. R., Holmer, L. E., Scleritome construction, biofacies, biostratigraphy and systematics of the tommotiid Eccentrotheca helenia sp. nov. from the Early Cambrian of South Australia. Palaeontology 54, 253–286 (2011).

8. Taylor, T. N., Taylor, E. L., Krings, M., Paleobotany: The Biology and Evolution of Fossil Plants (Academic Press, Amsterdam, Boston, Heidelberg, London, New York, Oxford, Paris, San Diego, San Francisco, Singapore, Sydney, Tokyo, ed. 2, 2009).

9. Berger, S., Kaever, M. J., Dasycladales: An Illustrated Monograph of a Fascinating Algal Order (Georg Thieme Verlag, Stuttgart, New York, 1992).

10. Gandin, A., Debrenne, F., Distribution of the archaeocyath-calcimicrobial bioconstructions on the Early Cambrian shelves. Palaeoworld 19, 222–241 (2010).

11. Debrenne, F., Rozanov, A., Zhuravle, A. Yu., Regular Archaeocyaths: Morphology, Systematic, Biostratigraphy, Palaeogeography, Biological Affinities (Centre National de la Recherche Scientifique, Paris, 1990).

12. Debrenne, F., Zhuravlev, A. Yu., Kruse, P. D., “Systematic descriptions: Archaeocyatha” in Treatise on Invertebrate Paleontology, Part E, Porifera (revised), v. 4-5 (Hypercalcified Porifera), P. A. Selden, Ed. (The University of Kansas Paleontological Institute, Lawrence, Kansas, 2015), pp. 923–1084.

13. Debrenne, F., Zhuravlev, A. Y., Kruse, P. D., “General features of Archaeocyatha” in Treatise on Invertebrate Paleontology, Part E, Porifera (Revised), Hypercalcified Porifera, Volume 4-5, P. A. Selden, Ed. (The University of Kansas Paleontological Institute, Lawrence, Kansas, 2015), pp. 845–922.

14. Korde, K. B., Novye predstaviteli sifonnikovykh vodoroslej [New representatives of the siphonous algae]. Materialy k Osnovam paleontologii 1, 67–75 (1957).

15. Korde, K. B., Vodorosli kembrija jugo-vostochnoj Sibirskoj platformy [Algae of the Cambrian from the South-Eastern Siberian platform]. Trudy Paleontologicheskogo Instituta Akademii Nauk SSSR 89, 1–147 (1961).

16. Riding, R., “Calcified algae and bacteria” in The ecology of the Cambrian radiation, Zhuravlev, A. Yu., Riding, R., Eds. (Columbia University Press, New York, 2001), pp. 445–473.

17. Wood, R. A., Zhuravlev, A. Yu., Debrenne, F., Functional biology and ecology of Archaeocyatha. Palaios 7, 131–156 (1992).

18. Flügel, E., “Fossils in Thin Section: It is Not That Difficult” in Microfacies of Carbonate Rocks: Analysis, Interpretation and Application (Springer, Berlin, Heidelberg, 2010), pp. 399–574.

19. Kenrick, P., Li, C.-S., An early, non-calcified, dasycladalean alga from the Lower Devonian of Yunnan Province, China. Rev. Palaeobot. Palynol. 100, 73–88 (1998).

20. Kenrick, P., Vinther, J., Chaetocladus gracilis n. sp., a non-calcified Dasycladales from the Upper Silurian of Sk åne, Sweden. Rev. Palaeobot. Palynol. 142, 153–160 (2006).

21. LoDuca, S. T., The green alga Chaetocladus (Dasycladales). J. Paleontol. 71, 940–949 (1997).

22. LoDuca, S. T., Kluessendorf, J., Mikulic, D. G., A new noncalcified Dasycladalean alga from the Silurian of Wisconsin. J. Paleontol. 77, 1152–1158 (2003).

23. LoDuca, S. T., Bykova, N., Wu Xiao, M., Zhao, S. Y., Seaweed morphology and ecology during the great animal diversification events of the early Paleozoic: A tale of two floras. Geobiology 15, 588–616 (2017).

24. Anderson, L. I., First non-calcified dasycladalean alga from the Carboniferous. N. Jb. Geol. Paläont. Abh. 251, 119–128 (2009).

25. Mastik, V., Tinn, O., New dasycladalean algal species from the Kalana Lagerstätte (Silurian, Estonia). J. Paleontol. 89, 262–268 (2015).

26. LoDuca, S. T., Tetreault, D. K., Ontogeny and reproductive functional morphology of the macroalga Wiartonella nodifera n. gen. n. sp. (Dasycladales, Chlorophyta) from the Silurian Eramosa Lagerstätte of Ontario, Canada. J. Paleontol. 91, 1–11 (2017).

27. Pettersson, J., Ahlberg, P., Lindskod, A., Lindgren, J., Eriksson, M. E., The fossil alga Chaetocladus gracilis revisited: New material from the Silurian of Sweden. GFF 142, 304–308 (2020).

28. Kruse, P. D., Debrenne, F., Ajax Mine archaeocyaths: A provisional biozonation for the upper Hawker Group (Cambrian stages 3–4), Flinders Ranges, South Australia. Australasian Palaeontological Memoirs 53, 1–238 (2020).

29. Yang, B., Steiner, M., Keupp, H., Early Cambrian palaeobiogeography of the Zhenba– Fangxian Block (South China): Independent terrane or part of the Yangtze Platform? Gondwana Research 28, 1543–1565 (2015).

30. Rowland, S. M., Shapiro, R. S., “Reef patterns and environmental influences in the Cambrian and Earliest Ordovician” in Phanerozoic Reef Patterns, Kiessling, W., Flügel, E., Golonka, J., Eds. (SEPM (Society of Sedimentary Geology), Tulsa, Oklahoma, USA, 2002), SEPM Specian Publication, pp. 95–128.

31. Mount, J. F., Signor, P. W., “Faunas and facies - fact and artifact: Paleoenvironmental controls on the distribution of early Cambrian faunas” in Origin and Early Evolution of the Metazoa, Lipps, J. H., Signor, P. W., Eds. (Springer Science and Business Media New York, New York, 1992), pp. 27–51.

32. Debrenne, F., Gandin, A., Rowland, S. M., Lower Cambrian bioconstruction in northwestern Mexico (Sonora). Depositional setting, paleoecology and systematics of archaeocyaths. Geobios 22, 137–195 (1989).

33. Debrenne, F., Rozanov, A., Paleogeographic and stratigraphicdistribution of Regular Archaeocyatha (Lower Cambrian fossils). Geobios 16, 727–736 (1983).

34. Garcia-Porta, J., Sol, D., Pennell, M., Sayol, F., Kaliontzopoulou, A., Botero, C. A., Niche expansion and adaptive divergence in the global radiation of crows and ravens. Nat Commun. 13, 2086 (2022).

35. Luo, C., Yang, A. H., Zhuravlev, Reitner, A. Yu., J., Vauxiids as descendants of archaeocyaths: a hypothesis. Lethaia 54, 700–710 (2021).

