## Supplementary_materials for "Not dasycladalean alga, but an Odyssey of the earliest Phanerozoic animal reef-builders"

**The PDF file includes:**

Supplementary Fig. 1
Supplementary Fig. 2
Supplementary Table 1

**Other Supplementary Materials for this manuscript include the following:**

Supplementary Dataset 1

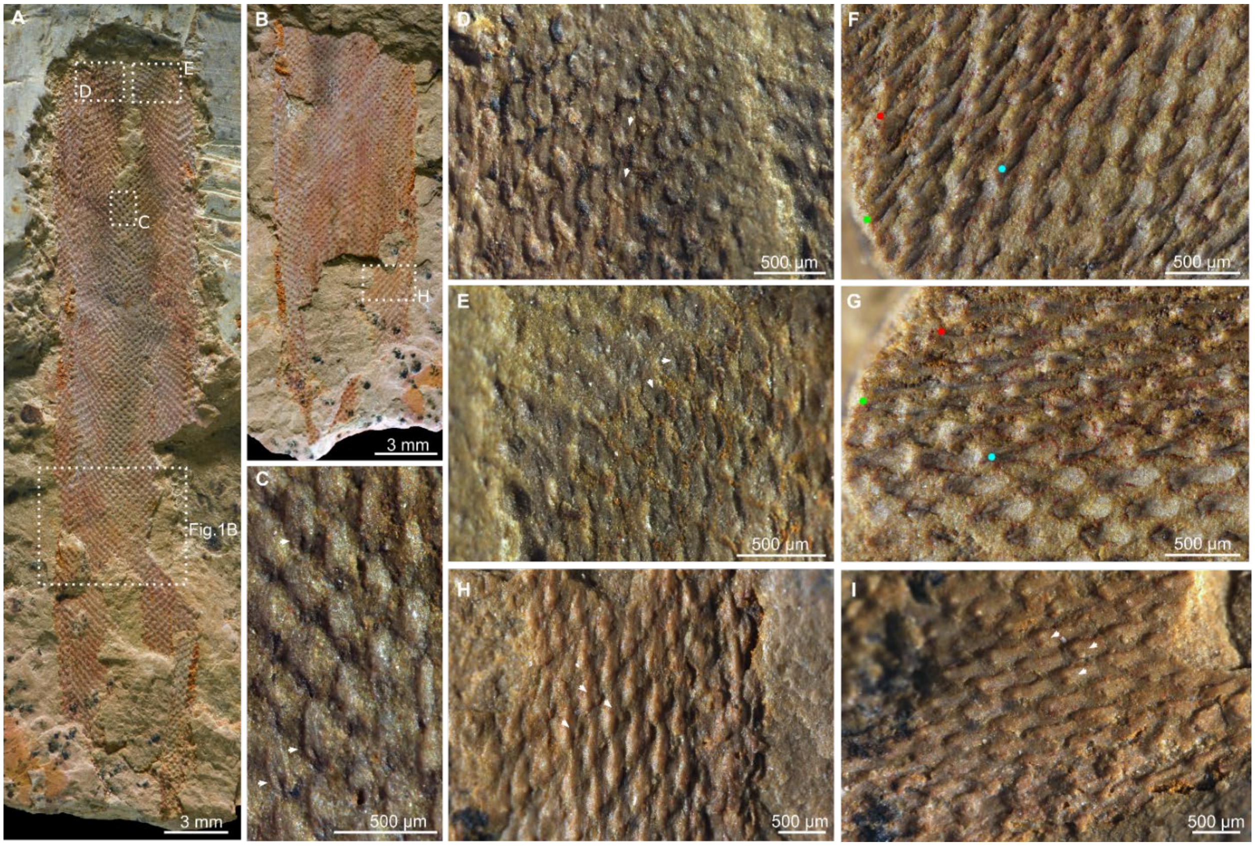


**Supplementary Fig. 1** Wall structures of possible archaeocyaths ASP1 from the Chengjiang Lagerstätte. **a, b** Counterparts of AC-02-JS, **a** is AC-02-JSa, **b** is AC-02-JSb. Areas in rectangles enlarged in **c–e, h** and Fig. 1b. **c–e** Close view of the inner surface, showing the presence of the tiny pores (arrows). **f, g** Part of AC-JS-08A in different angles of illumination, showing flattened canals forming rounded depressions on the surface. Dots indicate the same positions in the paired images. **h, i** Close view of the outer surface of part of AC-02-JSb, in different angles of illumination. Arrows indicate the tiny bulges beside the normal tubular canals, the provenance of which is uncertain.

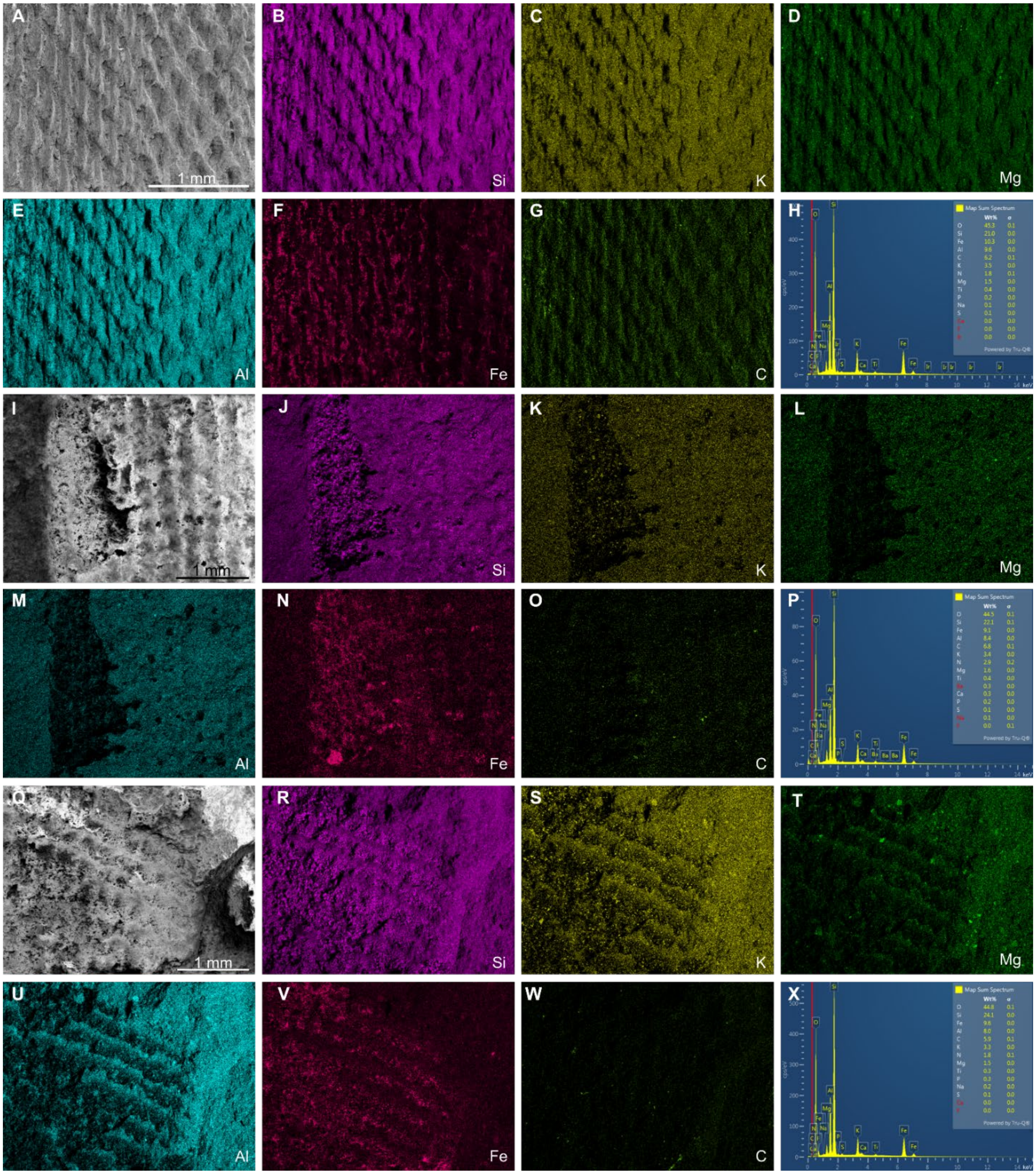


**Supplementary Fig. 2** Detailed elemental distribution and abundance of the investigated areas in the main text. **a–h** correspond to the elemental map in Fig. 1l. **i–p** correspond to the elemental map in Fig. 2c. **q–x** correspond to the elemental map in Fig. 2i.

**Supplementary Table 1 List of studied fossil specimens with brief descriptions.**

| Article No. | Structure arrangement | Notes (completeness; the preserved part of the organism) | Length (mm) | Width (mm) | Photos in the article |
|---|---|---|---|---|---|
| AC-02-JSa,b | diagonal | incomplete; top-middle part | 36.8 | 6.8 | Fig. 1b, c; Supplementary Fig. 1a–e, h, i |
| AC-03-JS | diagonal | almost complete, lack topmost part and holdfast | 48.5 | 8.9 | Fig. 1a, d, f–i |
| AC-04-JSa,b | diagonal | incomplete, middle-basal part | 62.8 | 7.2 | |
| AC-05-JSa,b | diagonal | incomplete, lack upper part | 48.7 | 6.8 | |
| AC-06-JSa,b,c | diagonal | incomplete, lack upper part | 27.4 | 4.0 | |
| AC-08-JSa,b | diagonal | incomplete; upper part | 11.9 | 6.6 | Fig. 1j–l |
| AC-07-SKa,b | orthogonal | incomplete; fragment (the exposed surface) | 7.3 | 6.2 | Fig. 2e–j |
| AC-09-SK | orthogonal | incomplete; upper to lower part | 34.5 | 6.8 | |
| AC-10-SKa | orthogonal | incomplete; middle part | 38.0 | 7.5 | Fig. 2a–d |
| AC-11-SK | orthogonal | complete | 85.5 | 9.5 | |
| AC-12-Ska,b | orthogonal | incomplete; lack basal part | 58.0 | 9.5 | |
| AC-13-SK | orthogonal | incomplete; lack basal part | 41.3 | 7.1 | |
| AC-14-SK | orthogonal | incomplete; fragment | 7.7 | 5.3 | |
| AC-15-SK | orthogonal | incomplete; lower-basal part | 35.3 | 4.6 | |
| AC-16-SK | orthogonal | incomplete; middle part | 51.3 | 8.8 | |
| AC-17-SK | orthogonal | incomplete; lower-basal part | 30.2 | 5.6 | |
| AC-18-SK | orthogonal | incomplete; middle part | 29.0 | 8.0 | |
| AC-19-SK | orthogonal | incomplete; middle part | 39.0 | 6.4 | |
| AC-20-SK | orthogonal | incomplete; middle part | 27.4 | 5.2 | |
| AC-21-SK | orthogonal | incomplete; top-upper part | 36.1 | 7.6 | |
| AC-22-JS | - | Thin section | | | |
| AC-23-JS | - | Thin section | | | |
| AC-24-SK | orthogonal | incomplete; upper part | 25.9 | 9.5 | |
| AC-25-SK | orthogonal | incomplete; lack basal part | 43.5 | 8.0 | |
| AC-26-SK | orthogonal | incomplete; lack basal part | 57.5 | 7.5 | |
| AC-27-SK | orthogonal | incomplete; middle part | 43.0 | 9.5 | |

**Supplementary Dataset 1. (in a separate file) Original data and parameters for the micro-CT scanning of the specimen illustrated in Fig. 2f–g.** Data are accessible at https://... (original data will be uploaded upon the acceptance of the manuscript).
